# Diversity of responses of soil saprobic fungi to recurring heat events

**DOI:** 10.1101/733923

**Authors:** Aleksandra Szymczak, Masahiro Ryo, Matthias C. Rillig

**Author notes:** Author for correspondence: M. Rillig.

## Abstract

As a consequence of ongoing climate change, the frequency of extreme heat events is expected to increase. Recurring heat pulses may disrupt functions supported by soil microorganisms, thus affecting the entire ecosystem. However, most perturbation experiments only test effects of single heat events, and therefore it remains largely unknown how soil microorganisms react to repeated pulse events. Here we present data from a lab experiment exposing 32 filamentous fungi, originally isolated from the same soil, to sequential heat perturbations. Soil saprobic fungi isolates were exposed to one or two heat pulses: mild (35°C/2h), strong (45°C/1h), or both in sequence (35°C/2h+45°C/1h), and we assessed growth rate. Out of the 32 isolates 13 isolates showed an antagonistic response, 3 isolates a synergistic response and 16 isolates responded in an additive manner. These differences in species responses to the thermal environment may contribute to species coexistence, and such dissimilarities in thermal perturbation responses may be a key aspect influencing ecosystem services that soil saprobic fungi support.

## Introduction

Climate warming is threatening ecosystems worldwide (IPCC 2018). Climate change does not only mean increased temperature averages but also increased frequency of extreme events, such as summer heatwaves (Bárcenas-Moreno et. al 2009; Hanson et. al 2006; Frank et al. 2015, IPCC 2018). Such extremes can profoundly influence individual physiological performance and fitness, phenotypic plasticity, demography and population dynamics, species interactions, and community structure (Vázquez et al. 2015 and references therein), probably even more so than an increase in mean conditions (Thompson et al. 2013).

Performances of soil microbes under an elevated average temperature have been widely investigated (Hortal et al. 2016), but the responses to heat pulse perturbations are understudied (Jentsch et al. 2007; Kreyling & Beier 2013). Temperature pulse perturbations occurring within a short period of time can be especially damaging, because soil organisms may not be able to adjust their physiological response fast enough (Alley et al. 2003; Fisher & Knutti 2015). Nevertheless, most temperature-related experimental designs have minimized temperature variability to solely focus on the effects of one average temperature (Thompson et al. 2013; Lloret et al. 2012). Understanding how heat pulse perturbations affect soil microbial performance is an important issue in soil ecology that could lead to a better understanding of aboveground and belowground community functioning.

In particular, the responses to multiple perturbations are far less understood but important, since multiple events may result in diverse response types because the effect size of a single event may depend on the antecedent event, known as ecological memory or carryover effect (Ryo et al. 2019). Considering the growing threat of recurrent heatwaves, it has been recently advocated that experiments aimed at investigating the impact of extreme weather events should consider that today’s extremes will become the normal fluctuations in the future, and experimental designs should exceed the level of severity that we currently observe to provide an insight into an organism’s responses to conditions harsher than those under which they evolved (Bahn et al. 2014; Kayler et al. 2015; Foster et al. 2016). Nevertheless, how soil saprobic fungi respond to recurrent temperature pulse perturbations is largely unknown.

The combination of multiple stressors (perturbations) can result in additive effects, detrimental effects (i.e. synergism) or cause a reduction in effects (i.e. antagonism) (Mittler 2006). Additive effects means that stressors do not interact and therefore the combined effect is simply the sum of each effect. Synergy results from a positive interaction, exceeding the sum of negative effects caused by each single stress event (Côté et al. 2016). Antagonism means that the combined effect is lower than the sum of each (negative) effect,such as observed in the form of stress priming ability (Rillig et al. 2015; Hilker et al. 2016). Priming ability means that a first exposure to a milder stress event induces protection mechanisms, consequently alleviating the effect of a subsequent stronger stress event (Rillig et al. 2015; Hilker et al. 2016; Andrade-Linares et al. 2016). While such different response types are theoretically possible, there is no study testing if such diverse responses to recurrent heat pulses are present in soil microbes co-occurring in the same environment. Additionally, studies on pulse temperature perturbations focus mostly on the community perspective, not providing information on species-level physiological responses (Norris et al. 2002; Allison & Martiny 2008; Crowther & Bradford 2013; Bérard et al.2015; Zhou et al. 2017).

The purpose of the present study was to investigate the diversity of physiological responses of soil filamentous fungi to sequential high-temperature pulses exceeding current adverse extreme conditions. We investigated how recurrent temperature pulses affect the performance of individual fungi from a set of 32 soil filamentous fungi that had been isolated from the same soil. We exposed fungi to one or two high-temperature pulse perturbations differing in magnitude (35°C/2h - mild (**M**), 45°C/1h - severe (**S**), and the sequence of these two perturbations 35°/2h+45°C/1h (**MS**)) and measured growth responses (colony extension rates). We expected the following: (1) exposure of soil saprobic fungi to recurrent temperature pulses will lead to diverse, isolate-specific responses, and (2) the diversity of responses to recurrent pulse temperature disturbance is phylogenetically conserved.

## Materials and methods

### Isolates

Isolates of 32 soil fungi (Supplementary materials, Table S1) were originally cultured from the top 10 cm of soil in a semi-arid grassland in Mallnow Lebus, Brandenburg, Germany (Andrade-Linares et al. 2016). To obtain material for the experiment, 6.5 mm plugs were taken from the edge of fungal colonies and placed centrally on 9 cm-diameter Petri dishes with PDA medium. Plates were then incubated at 22°C for 1 to 5 days, depending on individual colony extension rates, to obtain fresh and actively growing material for inoculation. Then, fungi were re-inoculated on fresh PDA plates and placed in incubators for the experiment.

### Heat treatment

In the field where the fungi were collected, the topsoil (at approx. 10 cm depth) temperature recorded in the year 2018 (52°52.778’N, 14°29.349’E) (Andrade-Linares et al. 2016) reached 32°C (Andrade-Linares et al. 2016, Dr. Max-Bernhard Ballhausen - personal information, data not shown). We used 35°C/2h as the mild perturbation (**M**) pulse temperature, since it is a temperature outside of the range of optimal growth conditions for half of the tested isolates, and it resulted in growth reduction in half of the tested fungi (Andrade-Linares et al. 2016). As the severe perturbation (**S**), we used 45°C applied for 1h since the responses to this temperature were severe for most of the isolates in our set (Andrade-Linares et al. 2016).

The full factorial experiment consisted of the following treatments: control (**C**) 22°C; mild perturbation (**M**) (35°C/2h); severe perturbation (**S**) (45°C/1h); sequence of the two perturbations (**MS**) (35°C/2h+45°C/1h).

Temperature pulses were applied uniformly to all 32 soil saprobic fungi. Each treatment consists of three replicates, with incubators used as experimental units. First, samples were incubated for 2-6 days to allow a fungal colony to begin growing from the inoculated plug. Then fungi were exposed to the different pulse temperature perturbations.

### Measurements

#### Trait measurements

The colony diameter of each isolate was measured for each Petri dish in two directions, at right angles to each other. Such measurements were taken four times – the first time before starting heat treatments to determine initial colony size, and then three more times after the treatment to define the response to heat exposure. The frequency of diameter measurements was isolate dependent and taken daily for fast-growing fungi or every 2-4 days for slow-growing individuals.

Thereby, in total 1,536 data points were acquired (i.e. 32 isolates × (2 temperatures magnitudes (M,S) + 1 temperature combination (MS) + control (C)) × 3 replicates × 4 time points). The diameter was measured repeatedly to calculate colony extension rate (mm day^-1^)

### Statistical analyses

Colony extension rates after heat treatments were used as a response variable to applied temperature pulses. Treatment effects were tested with two-way ANOVA where factors were the applied temperature regimes: mild perturbation (M, yes/no) and stronger perturbation (S, yes/no). Note that the no-no combination indicates control, while the yes-yes combination indicates two perturbations. The significance level α was set to 0.05 with Benjamini-Hochberg correction, to control for experiment-wise type I error rate.

We used this analysis to classify responses of fungi. Specifically, our response classification is based on additive null model expectations, used to identify interactions (antagony, synergy, additivity) between multiple perturbations (Crain et al. 2008; Côté et al. 2016). The additive null model has been reported to fit responses such as growth (colony extension) of an organism and is consistent with the use of ANOVA for factorial experimental data (Piggott et al. 2015; Côté et al. 2016). Response types were assigned to three groups (see Table 1). The effect direction of the two single perturbation effects in this study could be double negative (both single perturbations reduce the growth rate of the fungal isolate), opposing (one single perturbation increases the growth rate and the other single perturbation decreases the growth rate of the fungal isolate) or double positive (both single perturbations increase the growth rate of the fungal isolate) (Fig. 1). Those effect directions are crucial to assign response to interaction types (Crain et al. 2008).

**Figure 1.**
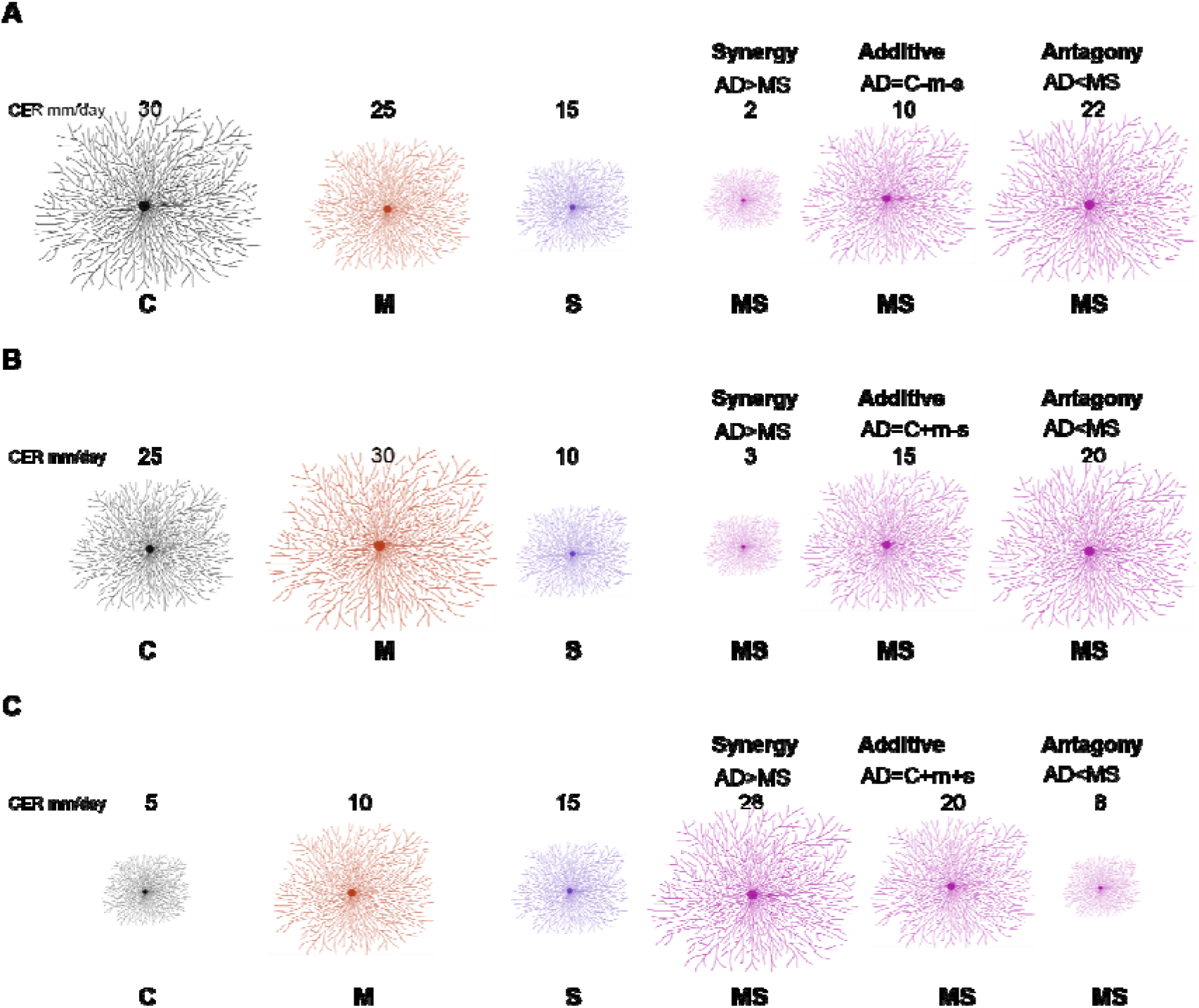
Redrawn from Crain et al. (2008) with adjusted conceptual approach for interpreting interaction types from factorial experiment response data. A full factorial study includes the following treatments: control (C), mild perturbation (M), strong perturbation (S) and both (MS). The three panels illustrate different combinations of individual responses: double negative (A), opposing (B), double positive (C). The numbers indicate example values of colony extension rate (proportional to the size of the mycelium). Interaction types (synergy, additive, antagony) depend on MS response in comparison with the sum (AD) of individual responses of single perturbations (m=C-M; s=C-S).

**Table 1.** Interaction types describing the outcome of multiple stressors (perturbations) following Crain et al. (2008); Côté et al. (2016). M (milder perturbation), S (stronger perturbation), MS (both perturbations applied).

|  | Individual treatment response |  |  |
| --- | --- | --- | --- |
|  | Additive | Non-additive |  |
|  |  | Synergy | Antagony |
| Condition | No significant interaction term between milder perturbation (M) and stronger perturbation (S) | A significant interaction term between milder perturbation (M) and stronger perturbation (S) | A significant interaction term between milder perturbation (M) and stronger perturbation (S) |
| Outcome category | The effect of MS is the equivalent of the addition of the single effects of M and S | The effect of MS is stronger than the added effects of M and S | The effect of MS is weaker than the added effects of M and S |

In addition, our study can be viewed in the context of stress priming, for which criteria were previously established (Andrade-Linares et al. 2016): (1) negative effect of strong perturbation (S); (2) significant interaction between mild and strong perturbations, and; (3) the interaction term has a positive sign (MS>S).

## Results

Fungal isolates showed a range of responses to the applied sequences of temperature perturbation (Fig. 2, Fig. 3). There was a significant interaction term for the two perturbations (M:S) in 16 tested isolates, and these were further categorized as antagonistic (13 isolates) or synergistic (3 isolates; Table 2). Isolates that did not meet the criterion of a significant interaction term (M:S) were assigned to the category ‘additive’ (16 isolates) (Tables 1 and 2).

**Figure 2.**
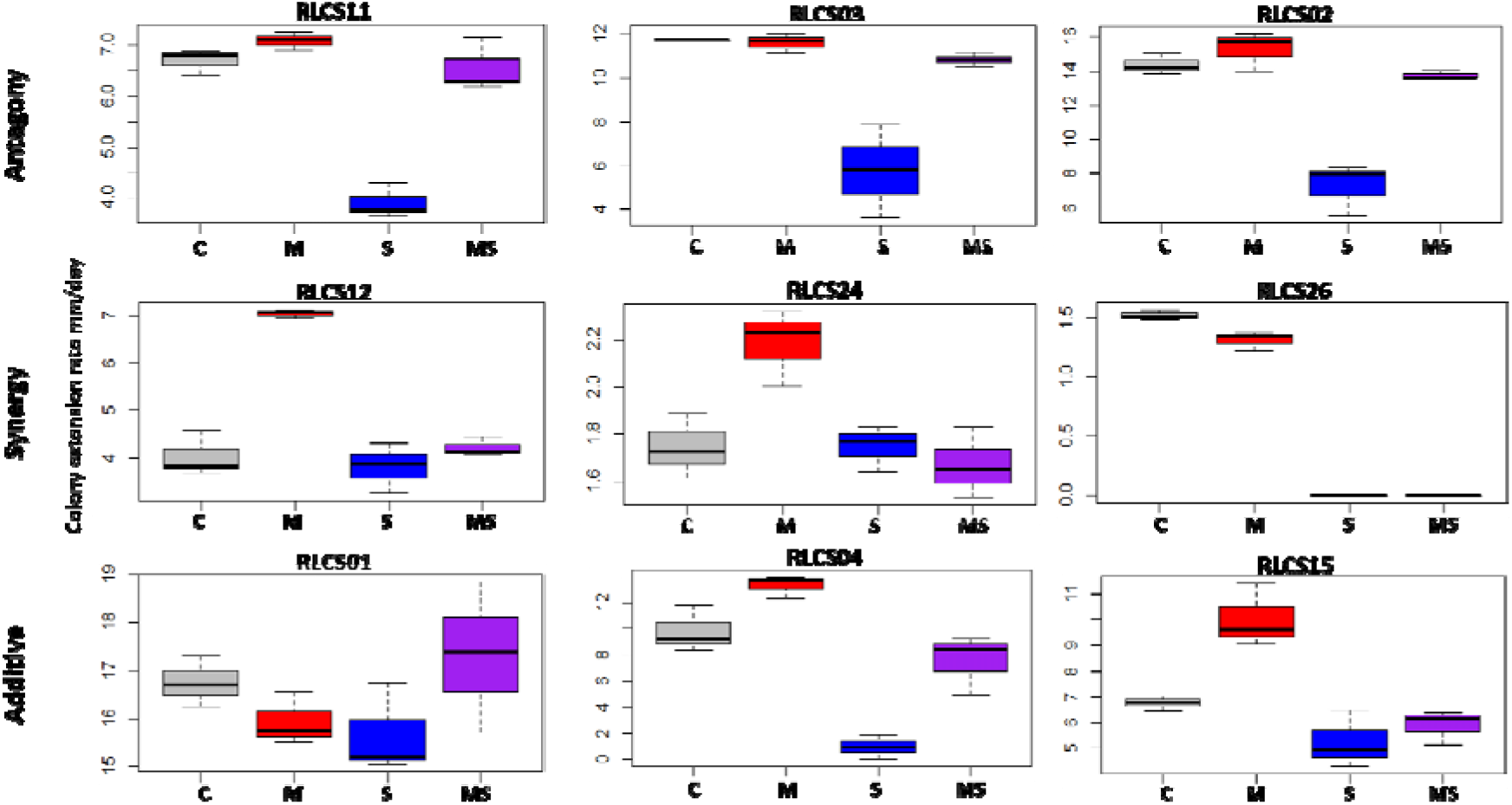
Examples of response categories (based on colony extension rate) to applied recurrent heat pulse perturbations: antagony (shown are 3 of the 13 isolates so categorized), synergy (all 3 isolates in this category are shown), additive (3 of the 16 isolates so categorized are depicted). For a full figure containing data for all 32 isolates see Supplementary materials (Figure S1, Supplementary materials).

**Figure 3.**
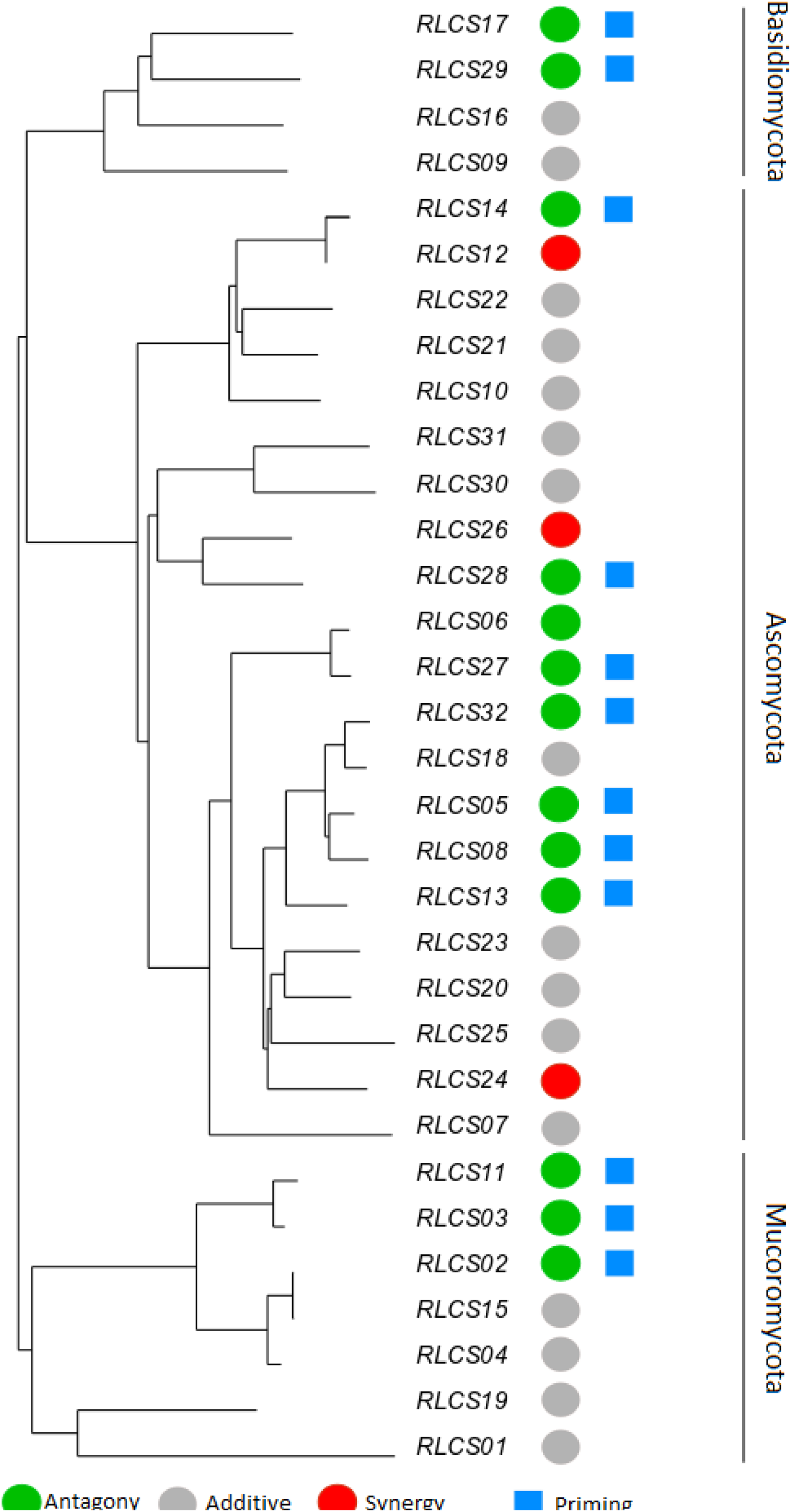
Phylogenetic relationship of a set of soil fungal isolates originating from the same field site with the representation of diverse responses of fungal colony extension rate to recurrent heat pulse perturbations (2h mild perturbation 35°C + 1h strong perturbation 45°C). The colored circles represent response types: antagony (green) (13 isolates); synergy (red) (3 isolates); additive (grey) (16 isolates). Blue squares mark isolates (12) that showed priming ability (based on criteria of Andrade-Linares et al. (2016)). The remaining isolates (19) did not meet the criteria of priming ability. The neighbor-joining tree was based on ITS and LSU regions; detailed information on isolates is in Supplementary materials (Supplementary materials, Table S1).

**Table 2.**
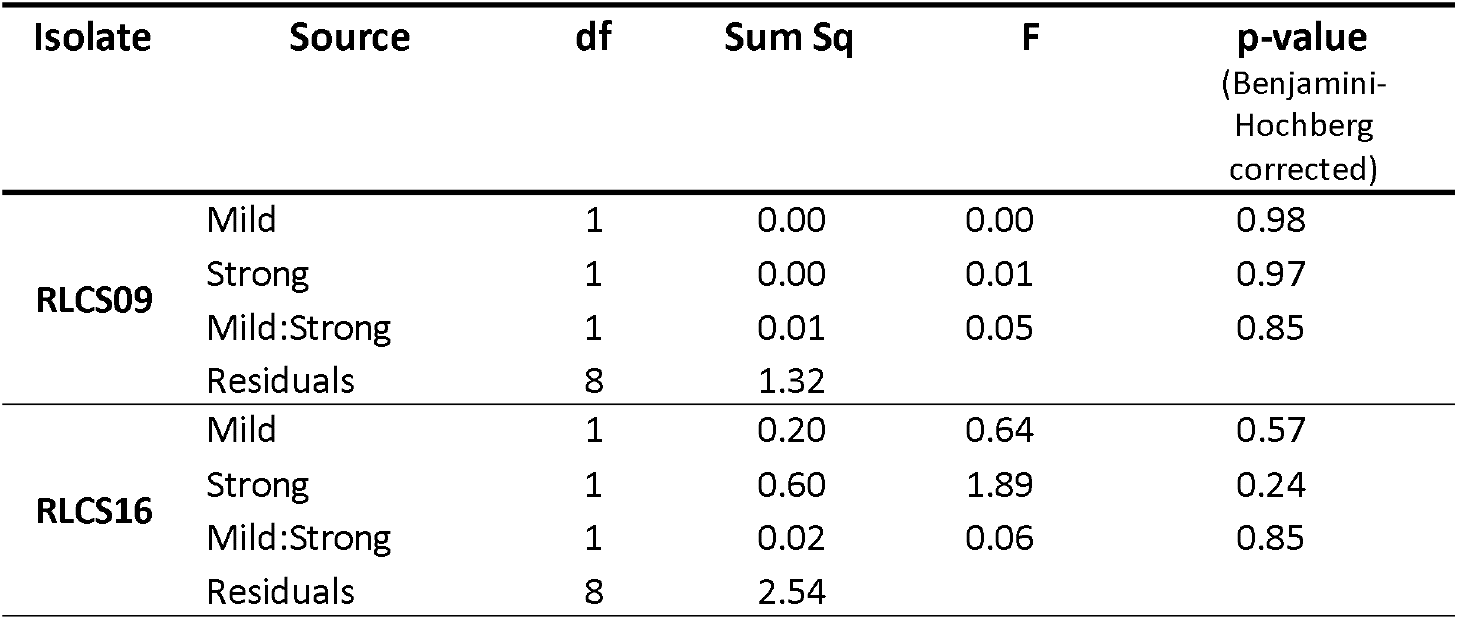
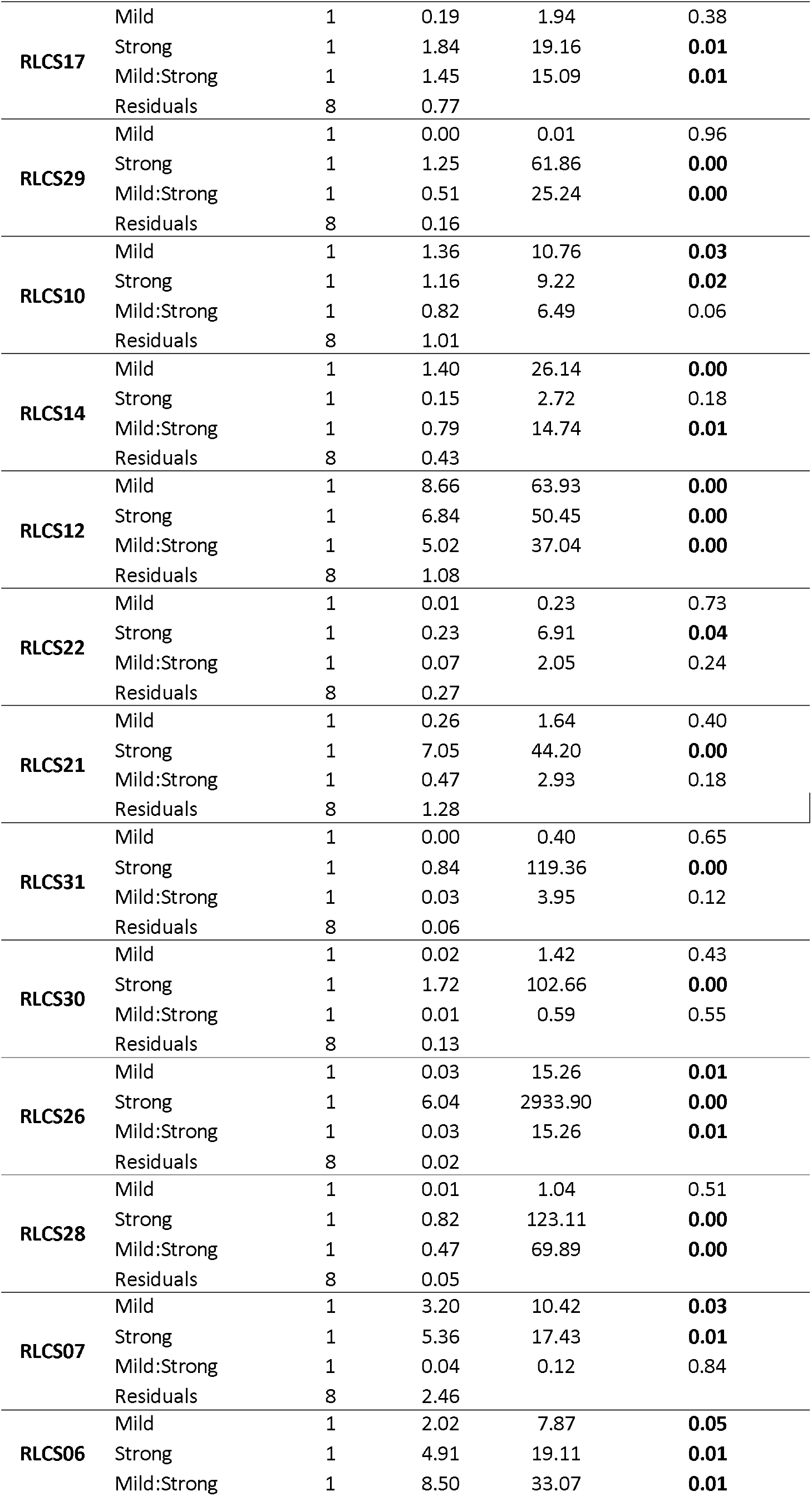
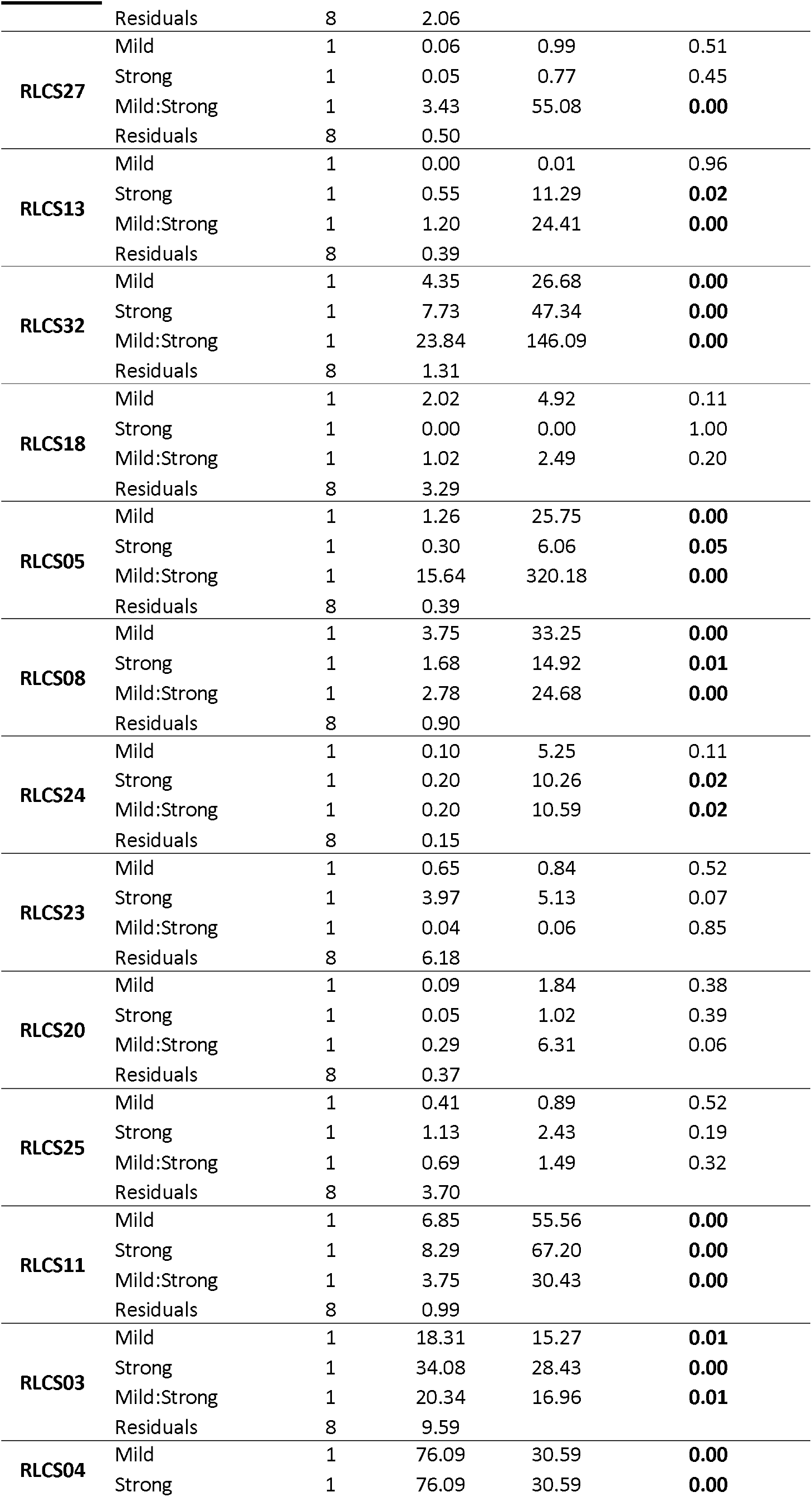
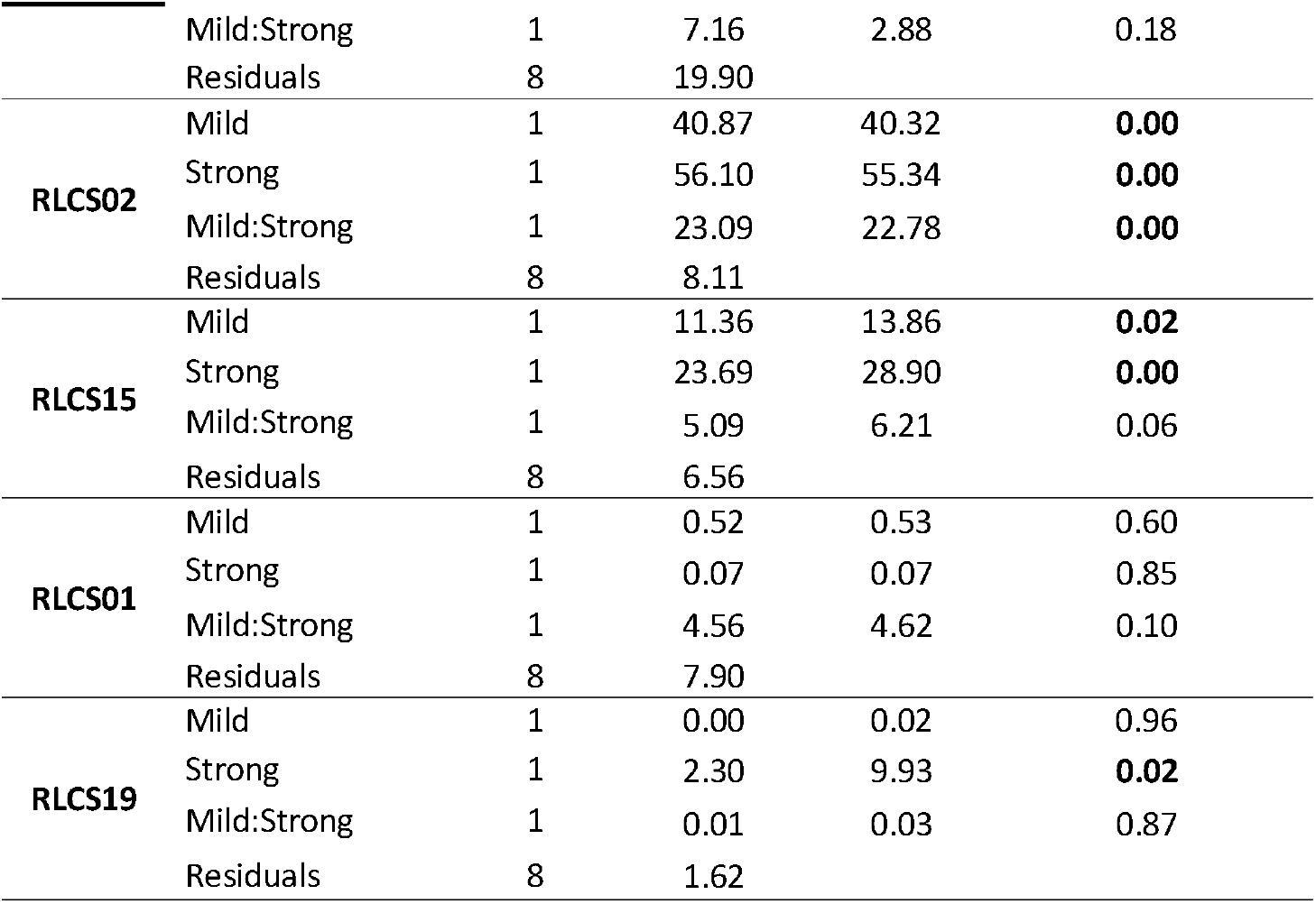
Analysis of variance results of effects of mild (M, yes/no), strong (S, yes/no) and both heat stress events (MS). The full results table is available in Supplementary materials (Supplementary materials, Table S2).

The observed antagonistic response is largely congruent (with small differences in categorization occurring, since two different statistical approaches are used - one isolate was classified differently) with what defines the priming ability of an organism (Andrade-Linares et al. 2016). We found priming responses in 12 out of the 32 isolates – 3 Mucoromycotina, 2 Basidiomycota and 7 Ascomycota. That is, for these 12 isolates, growth was enhanced when fungi were exposed to a mild temperature pulse (M) before the severe temperature (MS), compared to when they were only experiencing the severe pulse (S). For the remaining isolates exposure to experimental conditions of sequential perturbation (MS) did not lead to increased performance.

## Discussion

Here we show for the first time that exposure of soil saprobic fungi isolates to the same sequence of pulse temperature perturbations results in a range of response types that include additive effect, antagony and synergy. The applied temperature pulse regime was exceeding the temperatures that are currently typically observed at the site from which isolates were obtained. The biggest proportion of isolates (16/32) responded in an additive manner to the perturbations. The synergistic response (two perturbations leading to decreased performance) was observed for 3/32 isolates, meaning that such a sequence of perturbations could limit rates of processes (e.g. decomposition) these species carry out. On the other hand, the same temperature pulse regime resulted in an antagonistic response (two perturbations leading to an increased performance of an organism) for over 40% of isolates.

The observed antagonistic response of 12 isolates is congruent with priming ability of isolates. Priming ability of 8 filamentous fungi isolates exposed to a radically different temperature sequence (5h/35°C, then 40°C/10h) has been shown previously (Andrade-Linares et al. 2016). The fact that a large proportion of our fungi showed this improved response to sequential stresses, even under different temperature regimes, may indicate that this priming ability is not an exceptional response, but rather a well-established phenomenon that can help fungi deal with adverse effects.

In addition, the observed exacerbation of growth inhibition of three treated isolates (RLCS06, RLCS12, RLCS24) due to sequential heat exposure has not been observed before. For these fungi, the first heat stress event clearly did not help them in dealing with the second, more severe heat pulse. These would therefore be interesting targets to study further in the context of climate change and heat extremes.

At the species level, physiological stress regimes are known to set biogeographic limits and determine microhabitat preferences. Organismal responses to extreme heat events include redirecting resources from growth to survival that may include transition to a dormancy state or sporulation. These variations in heat responses are species specific and may be caused by differences in cellular HSP (heat shock proteins) production, altered membrane composition and carbohydrate flux (Morano et al. 2012; Bernard et al. 2015). The difference in response to sequential temperature pulse perturbation in isolates originating from one fungal assembly may indicate differences in sensitivity and diverse stress tolerances.

Our results show that recurrent environmental perturbations such as extreme temperature events influence a group of soil filamentous fungi originating from the same site in various ways. Thus, the patterns of responses that they exhibit to the sequence of thermal pulses might be one of the factors that contribute to shaping soil fungal community composition. Such differences in the ‘thermal niche’ may contribute to coexistence of fungi in the community, much likely differences among species in other abiotic factors or resource utilization patterns.

## Conclusion

We focused on physiological responses of multiple isolates of soil filamentous fungi, originating from the same grassland to a sequence of thermal pulses. These fungal isolates revealed the full range of possible responses. This could have consequences for soil-borne ecosystem services, highlighting the potential importance of fungal biodiversity in maintaining such services, particularly in the context of climate change.

## Acknowledgement

We acknowledge funding for CRC 973 Project A1 ‘Priming and memory of stress organismic responses to stress’ from the German Research Foundation (DFG).

