## Supplementary material for "Diversity of responses of soil saprobic fungi to recurring heat events"

Table S1. Taxonomic identification of the soil filamentous fungi used in this study. The order of species in the table is by phylogeny. The neighbor-joining tree was based on the ITS (intergenic transcribed spacer) and a part of the large rRNA subunit (LSU). Phylogenetic annotations were based on bootstrap analysis, and assumed valid when supported in 80% of the bootstraps.

| **Strain ID** | **Genus species** | **Order** | **Phylum** | **NCBI Accession number** | **DSMZ** |
| --- | --- | --- | --- | --- | --- |
| RLCS09 | *Trametes versicolor* | Polyporales | Basidiomycota | KT582071 | DSM 100406 |
| RLCS16 | *Pleurotus sapidus* | Agaricales | Basidiomycota | KT582080 | DSM 100408 |
| RLCS17 | *Clitopilus sp.* | Agaricales | Basidiomycota | KT582089 | DSM 100324 |
| RLCS29 | *Macrolepiota excoriata* | Agaricales | Basidiomycota | KT582069 | DSM 100288 |
| RLCS10 | *Alternaria alternata* | Pleosporales | Ascomycota | KT582078 | DSM 100286 |
| RLCS14 | *Didymellaceae*  strain 2 | Pleosporales | Ascomycota | KT582077 | DSM 100404 |
| RLCS12 | *Didymellaceae*  strain 1 | Pleosporales | Ascomycota | KT582079 | DSM 100405 |
| RLCS22 | *Paraphoma chrysanthemicola* | Pleosporales | Ascomycota | KT582091 | DSM 100401 |
| RLCS21 | *Pyrenochaetopsis leptospora* | Pleosporales | Ascomycota | KT582065 | DSM 100327 |
| RLCS31 | *Cyphellophora sp.* | Chaetothyriales | Ascomycota | KT582074 | DSM 100328 |
| RLCS30 | *Exophiala equina* | Chaetothyriales | Ascomycota | KT582075 | DSM 100291 |
| RLCS26 | *Tetracladium marchalianum* | Helotiales | Ascomycota | KT582084 | DSM 100330 |
| RLCS28 | *Tricladium sp.* | Helotiales | Ascomycota | KT582085 | DSM 100323 |
| RLCS07 | *Amphisphaeriaceae*  strain 1 | Xylariales | Ascomycota | KT582088 | DSM 100284 |
| RLCS06 | *Chaetomium angustispirale* | Sordariales | Ascomycota | KT582096 | DSM 100400 |
| RLCS27 | *Thielavia inaequalis* | Sordariales | Ascomycota | KT582086 | DSM 100326 |
| RLCS13 | *Fusarium solani* | Hypocreales | Ascomycota | KT582073 | DSM 100290 |
| RLCS32 | *Fusarium oxysporum* | Hypocreales | Ascomycota | KT582095 | DSM 100409 |
| RLCS18 | *Gibberella sp.* | Hypocreales | Ascomycota | KT582068 | DSM 100287 |
| RLCS05 | *Fusarium sp.* | Hypocreales | Ascomycota | KT582097 | DSM 100403 |
| RLCS08 | *Gibberella tricincta* | Hypocreales | Ascomycota | KT582087 | DSM 100325 |
| RLCS24 | *Metarhizium marquandii* | Hypocreales | Ascomycota | KT582066 | DSM 100410 |
| RLCS23 | *Stachybotryaceae*  strain 1 | Hypocreales | Ascomycota | KT582090 | DSM 101519 |
| RLCS20 | *Purpureocillium lilacinum* | Hypocreales | Ascomycota | KT582081 | DSM 100329 |
| RLCS25 | *Hydropisphaera* sp. | Hypocreales | Ascomycota | KT582083 | DSM 100292 |
| RLCS11 | *Mortierella alpina*  strain 2 | Mortierellales | Mucoromycota | KT582070 | DSM 100289 |
| RLCS03 | *Mortierella alpina*  strain 1 | Mortierellales | Mucoromycota | KT582067 | DSM 100285 |
| RLCS04 | *Mortierella exigua* | Mortierellales | Mucoromycota | KT582094 | DSM 100322 |
| RLCS02 | *Mortierella elongata*  strain 1 | Mortierellales | Mucoromycota | KT582072 | DSM 100407 |
| RLCS15 | *Mortierella elongata*  strain 2 | Mortierellales | Mucoromycota | KT582092 | DSM 100402 |
| RLCS01 | *Mucor fragilis* | Mucorales | Mucoromycota | KT582076 | DSM 100293 |
| RLCS19 | *Umbelopsis isabellina* | Umbelopsidales | Mucoromycota | KT582093 | DSM 100331 |

Table S2. Analysis of variance (full result table) - effects of mild (M, yes/no), strong (S, yes/no) and both heat stress events (MS).

| **Isolate** | **Source** | **df** | **Sum Sq** | **Mean Sq** | **F** | **p-value** | **p-value** (Benjamini- Hochberg corrected) |
| --- | --- | --- | --- | --- | --- | --- | --- |
| RLCS09 | Mild | 1 | 0.0001 | 0.0001 | 0.0010 | 0.9800 | 0.9804 |
|  | Strong | 1 | 0.0010 | 0.0010 | 0.0060 | 0.9410 | 0.9715 |
|  | Mild:Strong | 1 | 0.0086 | 0.0086 | 0.0520 | 0.8250 | 0.8515 |
|  | Residuals | 8 | 1.3180 | 0.1648 |  |  |  |
| RLCS16 | Mild | 1 | 0.2032 | 0.2032 | 0.6400 | 0.4470 | 0.5719 |
|  | Strong | 1 | 0.6006 | 0.6006 | 1.8910 | 0.2060 | 0.2445 |
|  | Mild:Strong | 1 | 0.0196 | 0.0196 | 0.0620 | 0.8100 | 0.8515 |
|  | Residuals | 8 | 2.5402 | 0.3175 |  |  |  |
| RLCS17 | Mild | 1 | 0.1859 | 0.1859 | 1.9410 | 0.2011 | 0.3762 |
|  | Strong | 1 | 1.8360 | 1.8360 | 19.1610 | 0.0024 | 0.0051 |
|  | Mild:Strong | 1 | 1.4457 | 1.4457 | 15.0890 | 0.0047 | 0.0106 |
|  | Residuals | 8 | 0.7665 | 0.0958 |  |  |  |
| RLCS29 | Mild | 1 | 0.0002 | 0.0002 | 0.0090 | 0.9284 | 0.9583 |
|  | Strong | 1 | 1.2452 | 1.2452 | 61.8610 | 0.0000 | 0.0002 |
|  | Mild:Strong | 1 | 0.5081 | 0.5081 | 25.2420 | 0.0010 | 0.0036 |
|  | Residuals | 8 | 0.1610 | 0.0201 |  |  |  |
| RLCS10 | Mild | 1 | 1.3588 | 1.3588 | 10.755 | 0.0112 | 0.0298 |
|  | Strong | 1 | 1.1644 | 1.1644 | 9.216 | 0.0162 | 0.0246 |
|  | Mild:Strong | 1 | 0.8195 | 0.8195 | 6.486 | 0.0343 | 0.0630 |
|  | Residuals | 8 | 1.0107 | 0.1263 |  |  |  |
| RLCS14 | Mild | 1 | 1.3975 | 1.3975 | 26.143 | 0.0009 | 0.0038 |
|  | Strong | 1 | 0.1453 | 0.1453 | 2.718 | 0.1379 | 0.1764 |
|  | Mild:Strong | 1 | 0.7877 | 0.7877 | 14.737 | 0.0050 | 0.0106 |
|  | Residuals | 8 | 0.4276 | 0.0535 |  |  |  |
| RLCS12 | Mild | 1 | 8.662 | 8.662 | 63.93 | 0.0000 | 0.0012 |
|  | Strong | 1 | 6.836 | 6.836 | 50.45 | 0.0001 | 0.0004 |
|  | Mild:Strong | 1 | 5.019 | 5.019 | 37.04 | 0.0003 | 0.0019 |
|  | Residuals | 8 | 1.084 | 0.135 |  |  |  |
| RLCS22 | Mild | 1 | 0.00792 | 0.00792 | 0.234 | 0.6413 | 0.7329 |
|  | Strong | 1 | 0.23339 | 0.23339 | 6.909 | 0.0302 | 0.0440 |
|  | Mild:Strong | 1 | 0.06915 | 0.06915 | 2.047 | 0.1904 | 0.2437 |
|  | Residuals | 8 | 0.27023 | 0.03378 |  |  |  |
| RLCS21 | Mild | 1 | 0.262 | 0.262 | 1.640 | 0.2362 | 0.3978 |
|  | Strong | 1 | 7.050 | 7.050 | 44.196 | 0.0002 | 0.0005 |
|  | Mild:Strong | 1 | 0.468 | 0.468 | 2.931 | 0.1253 | 0.1783 |
|  | Residuals | 8 | 1.276 | 0.160 |  |  |  |
| RLCS31 | Mild | 1 | 0.0028 | 0.0028 | 0.396 | 0.5470 | 0.6479 |
|  | Strong | 1 | 0.8361 | 0.8361 | 119.363 | 0.0000 | 0.0000 |
|  | Mild:Strong | 1 | 0.0277 | 0.0277 | 3.952 | 0.0820 | 0.1250 |
|  | Residuals | 8 | 0.0560 | 0.0070 |  |  |  |
| RLCS30 | Mild | 1 | 0.0237 | 0.0237 | 1.415 | 0.2680 | 0.4295 |
|  | Strong | 1 | 1.7176 | 1.7176 | 102.664 | 0.0000 | 0.0001 |
|  | Mild:Strong | 1 | 0.0098 | 0.0098 | 0.586 | 0.4660 | 0.5521 |
|  | Residuals | 8 | 0.1338 | 0.0167 |  |  |  |
| RLCS26 | Mild | 1 | 0.031 | 0.031 | 15.26 | 0.0045 | 0.0144 |
|  | Strong | 1 | 6.041 | 6.041 | 2933.9 | 0.0000 | 0.0000 |
|  | Mild:Strong | 1 | 0.031 | 0.031 | 15.26 | 0.0045 | 0.0106 |
|  | Residuals | 8 | 0.016 | 0.002 |  |  |  |
| RLCS28 | Mild | 1 | 0.0069 | 0.0069 | 1.041 | 0.3380 | 0.5079 |
|  | Strong | 1 | 0.8194 | 0.8194 | 123.113 | 0.0000 | 0.0000 |
|  | Mild:Strong | 1 | 0.4651 | 0.4651 | 69.885 | 0.0000 | 0.0003 |
|  | Residuals | 8 | 0.0532 | 0.0067 |  |  |  |
| RLCS07 | Mild | 1 | 3.202 | 3.202 | 10.420 | 0.0121 | 0.0298 |
|  | Strong | 1 | 5.357 | 5.357 | 17.434 | 0.0031 | 0.0062 |
|  | Mild:Strong | 1 | 0.037 | 0.037 | 0.121 | 0.7367 | 0.8419 |
|  | Residuals | 8 | 2.458 | 0.307 |  |  |  |
| RLCS06 | Mild | 1 | 2.021 | 2.021 | 7.868 | 0.0230 | 0.0526 |
|  | Strong | 1 | 4.909 | 4.909 | 19.109 | 0.0024 | 0.0051 |
|  | Mild:Strong | 1 | 8.495 | 8.495 | 33.068 | 0.0004 | 0.0023 |
|  | Residuals | 8 | 2.055 | 0.257 |  |  |  |
| RLCS27 | Mild | 1 | 0.062 | 0.062 | 0.989 | 0.3490 | 0.5079 |
|  | Strong | 1 | 0.048 | 0.048 | 0.772 | 0.4050 | 0.4471 |
|  | Mild:Strong | 1 | 3.431 | 3.431 | 55.078 | 0.0001 | 0.0006 |
|  | Residuals | 8 | 0.498 | 0.062 |  |  |  |
| RLCS13 | Mild | 1 | 0.0005 | 0.0005 | 0.011 | 0.9194 | 0.9583 |
|  | Strong | 1 | 0.5548 | 0.5548 | 11.288 | 0.0099 | 0.0177 |
|  | Mild:Strong | 1 | 1.1999 | 1.1999 | 24.413 | 0.0011 | 0.0036 |
|  | Residuals | 8 | 0.3932 | 0.0491 |  |  |  |
| RLCS32 | Mild | 1 | 4.354 | 4.354 | 26.68 | 0.0009 | 0.0038 |
|  | Strong | 1 | 7.726 | 7.726 | 47.34 | 0.0001 | 0.0004 |
|  | Mild:Strong | 1 | 23.841 | 23.841 | 146.09 | 0.0000 | 0.0000 |
|  | Residuals | 8 | 1.306 | 0.163 |  |  |  |
| RLCS18 | Mild | 1 | 2.020 | 2.0197 | 4.918 | 0.0574 | 0.1148 |
|  | Strong | 1 | 0.000 | 0.0000 | 0.000 | 0.9981 | 0.9981 |
|  | Mild:Strong | 1 | 1.024 | 1.0240 | 2.493 | 0.1530 | 0.2040 |
|  | Residuals | 8 | 3.286 | 0.4107 |  |  |  |
| RLCS05 | Mild | 1 | 1.258 | 1.258 | 25.752 | 0.0010 | 0.0038 |
|  | Strong | 1 | 0.296 | 0.296 | 6.063 | 0.0392 | 0.0545 |
|  | Mild:Strong | 1 | 15.640 | 15.640 | 320.178 | 0.0000 | 0.0000 |
|  | Residuals | 8 | 0.391 | 0.049 |  |  |  |
| RLCS08 | Mild | 1 | 3.749 | 3.749 | 33.25 | 0.0004 | 0.0034 |
|  | Strong | 1 | 1.682 | 1.682 | 14.92 | 0.0048 | 0.0090 |
|  | Mild:Strong | 1 | 2.783 | 2.783 | 24.68 | 0.0011 | 0.0036 |
|  | Residuals | 8 | 0.902 | 0.113 |  |  |  |
| RLCS24 | Mild | 1 | 0.09973 | 0.0997 | 5.246 | 0.0512 | 0.1093 |
|  | Strong | 1 | 0.19504 | 0.1950 | 10.259 | 0.0126 | 0.0211 |
|  | Mild:Strong | 1 | 0.20134 | 0.2013 | 10.590 | 0.0116 | 0.0233 |
|  | Residuals | 8 | 0.15210 | 0.0190 |  |  |  |
| RLCS23 | Mild | 1 | 0.648 | 0.648 | 0.839 | 0.3865 | 0.5154 |
|  | Strong | 1 | 3.965 | 3.965 | 5.129 | 0.0533 | 0.0711 |
|  | Mild:Strong | 1 | 0.043 | 0.043 | 0.055 | 0.8198 | 0.8515 |
|  | Residuals | 8 | 6.184 | 0.773 |  |  |  |
| RLCS20 | Mild | 1 | 0.0858 | 0.0858 | 1.844 | 0.2116 | 0.3762 |
|  | Strong | 1 | 0.0477 | 0.0477 | 1.024 | 0.3413 | 0.3900 |
|  | Mild:Strong | 1 | 0.2935 | 0.2935 | 6.305 | 0.0363 | 0.0630 |
|  | Residuals | 8 | 0.3724 | 0.0466 |  |  |  |
| RLCS25 | Mild | 1 | 0.413 | 0.4126 | 0.892 | 0.3730 | 0.5154 |
|  | Strong | 1 | 1.125 | 1.1255 | 2.433 | 0.1570 | 0.1937 |
|  | Mild:Strong | 1 | 0.689 | 0.6888 | 1.489 | 0.2570 | 0.3164 |
|  | Residuals | 8 | 3.700 | 0.4625 |  |  |  |
| RLCS11 | Mild | 1 | 6.853 | 6.853 | 55.56 | 0.0001 | 0.0012 |
|  | Strong | 1 | 8.289 | 8.289 | 67.20 | 0.0000 | 0.0002 |
|  | Mild:Strong | 1 | 3.754 | 3.754 | 30.43 | 0.0006 | 0.0026 |
|  | Residuals | 8 | 0.987 | 0.123 |  |  |  |
| RLCS03 | Mild | 1 | 18.31 | 18.31 | 15.27 | 0.0045 | 0.0144 |
|  | Strong | 1 | 34.08 | 34.08 | 28.43 | 0.0007 | 0.0017 |
|  | Mild:Strong | 1 | 20.34 | 20.34 | 16.96 | 0.0033 | 0.0089 |
|  | Residuals | 8 | 9.59 | 1.20 |  |  |  |
| RLCS04 | Mild | 1 | 76.09 | 76.09 | 30.594 | 0.0006 | 0.0035 |
|  | Strong | 1 | 76.09 | 76.09 | 30.594 | 0.0006 | 0.0002 |
|  | Mild:Strong | 1 | 7.16 | 7.16 | 2.879 | 0.1282 | 0.1783 |
|  | Residuals | 8 | 19.90 | 2.49 |  |  |  |
| RLCS02 | Mild | 1 | 40.87 | 40.87 | 40.32 | 0.0002 | 0.0024 |
|  | Strong | 1 | 56.10 | 56.10 | 55.34 | 0.0001 | 0.0003 |
|  | Mild:Strong | 1 | 23.09 | 23.09 | 22.78 | 0.0014 | 0.0041 |
|  | Residuals | 8 | 8.11 | 1.01 |  |  |  |
| RLCS15 | Mild | 1 | 11.360 | 11.360 | 13.858 | 0.0058 | 0.0170 |
|  | Strong | 1 | 23.689 | 23.689 | 28.896 | 0.0007 | 0.0017 |
|  | Mild:Strong | 1 | 5.092 | 5.092 | 6.211 | 0.0374 | 0.0630 |
|  | Residuals | 8 | 6.558 | 0.820 |  |  |  |
| RLCS01 | Mild | 1 | 0.521 | 0.521 | 0.527 | 0.4884 | 0.6011 |
|  | Strong | 1 | 0.067 | 0.067 | 0.068 | 0.8004 | 0.8537 |
|  | Mild:Strong | 1 | 4.563 | 4.563 | 4.621 | 0.0638 | 0.1021 |
|  | Residuals | 8 | 7.900 | 0.987 |  |  |  |
| RLCS19 | Mild | 1 | 0.0037 | 0.0037 | 0.016 | 0.9033 | 0.9583 |
|  | Strong | 1 | 2.2963 | 2.2963 | 9.933 | 0.0161 | 0.0246 |
|  | Mild:Strong | 1 | 0.0067 | 0.0067 | 0.029 | 0.8700 | 0.8700 |
|  | Residuals | 8 | 1.6183 | 0.2312 |  |  |  |

Figure S1. Response categories and full data for of all 32 fungal isolates (based on colony extension rate) to recurrent heat pulse perturbations: synergy, antagony, and additive response.

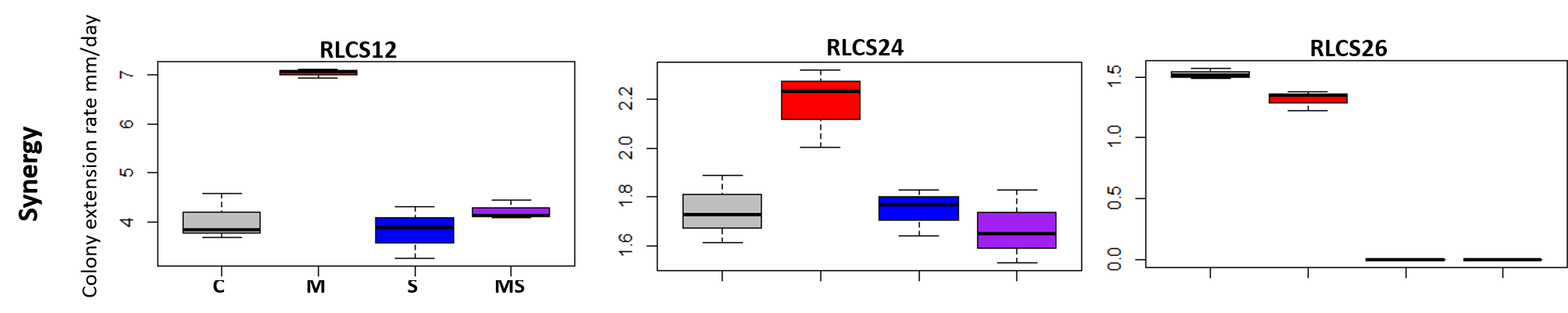

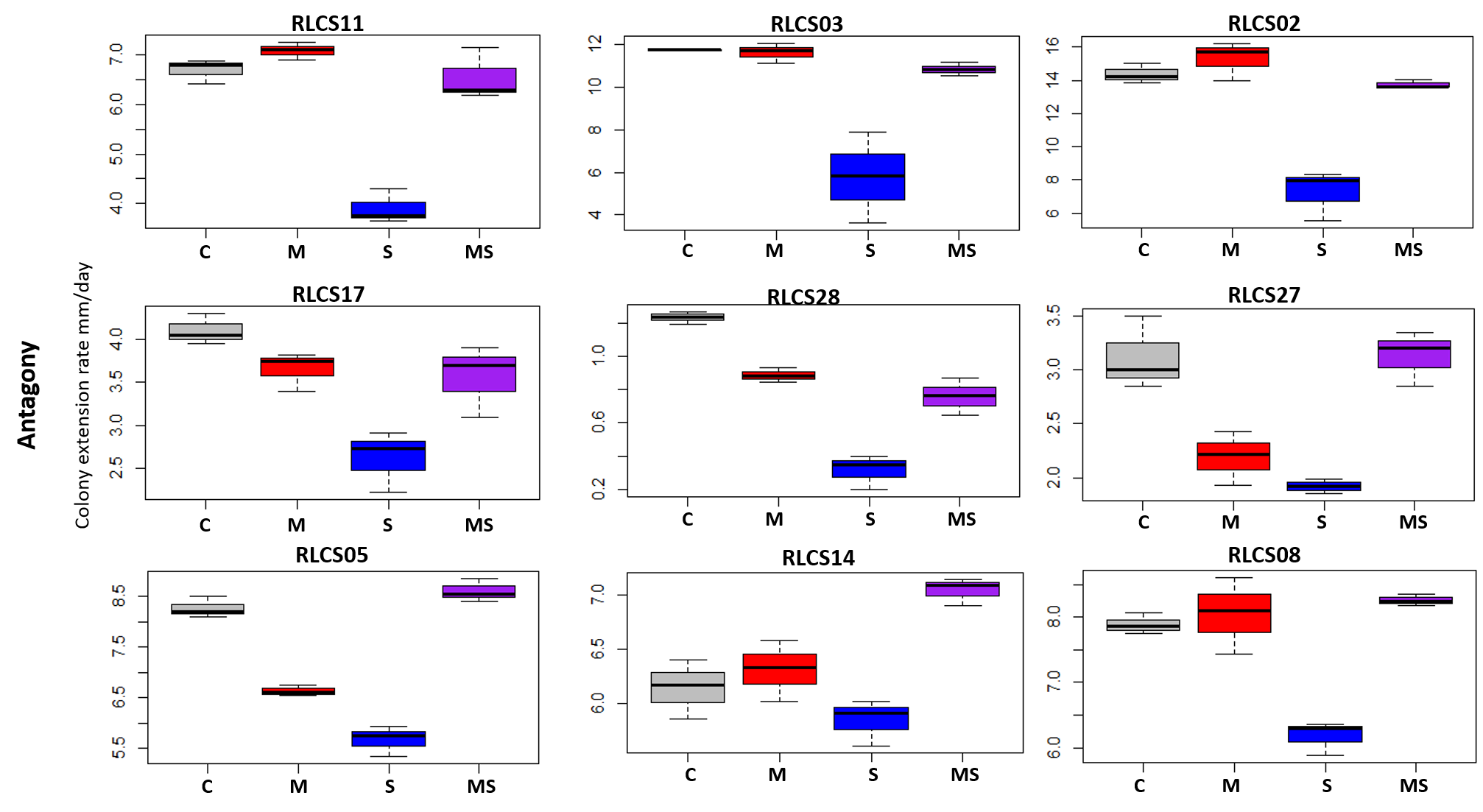

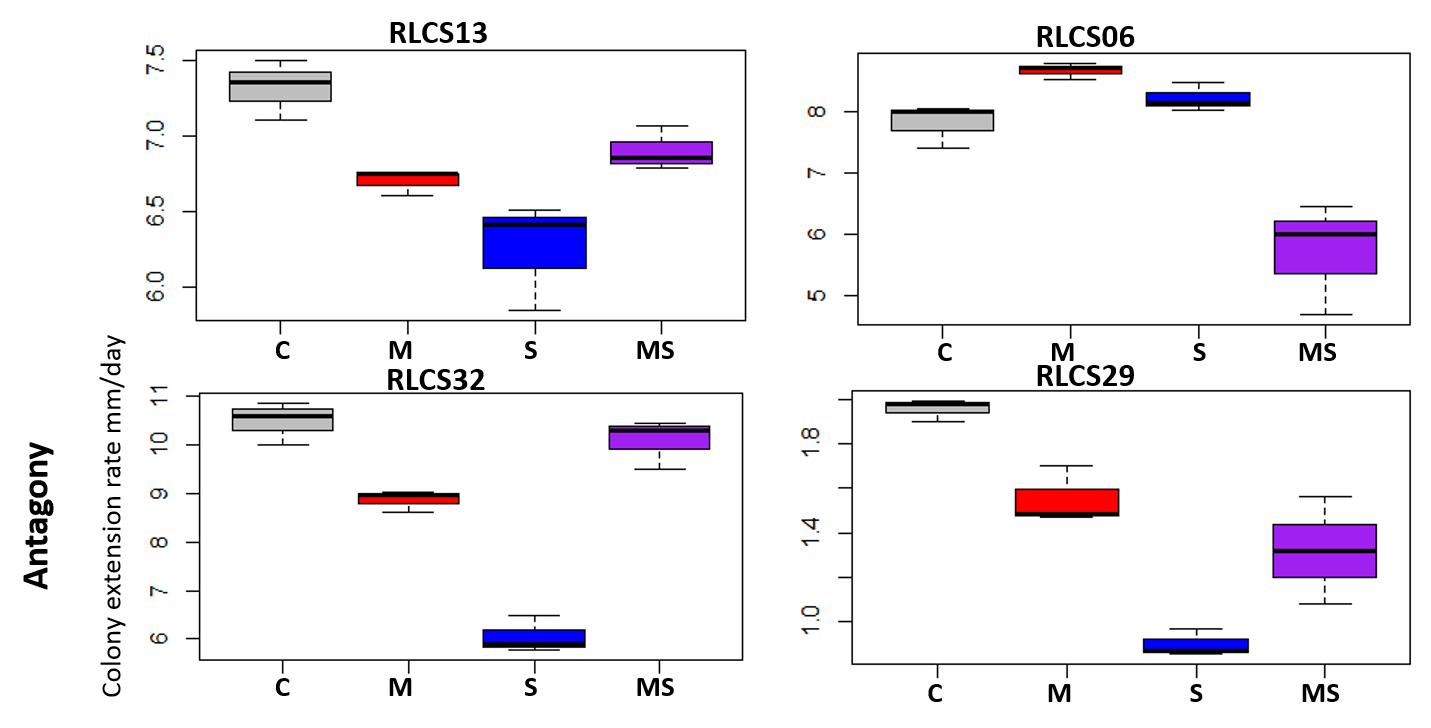

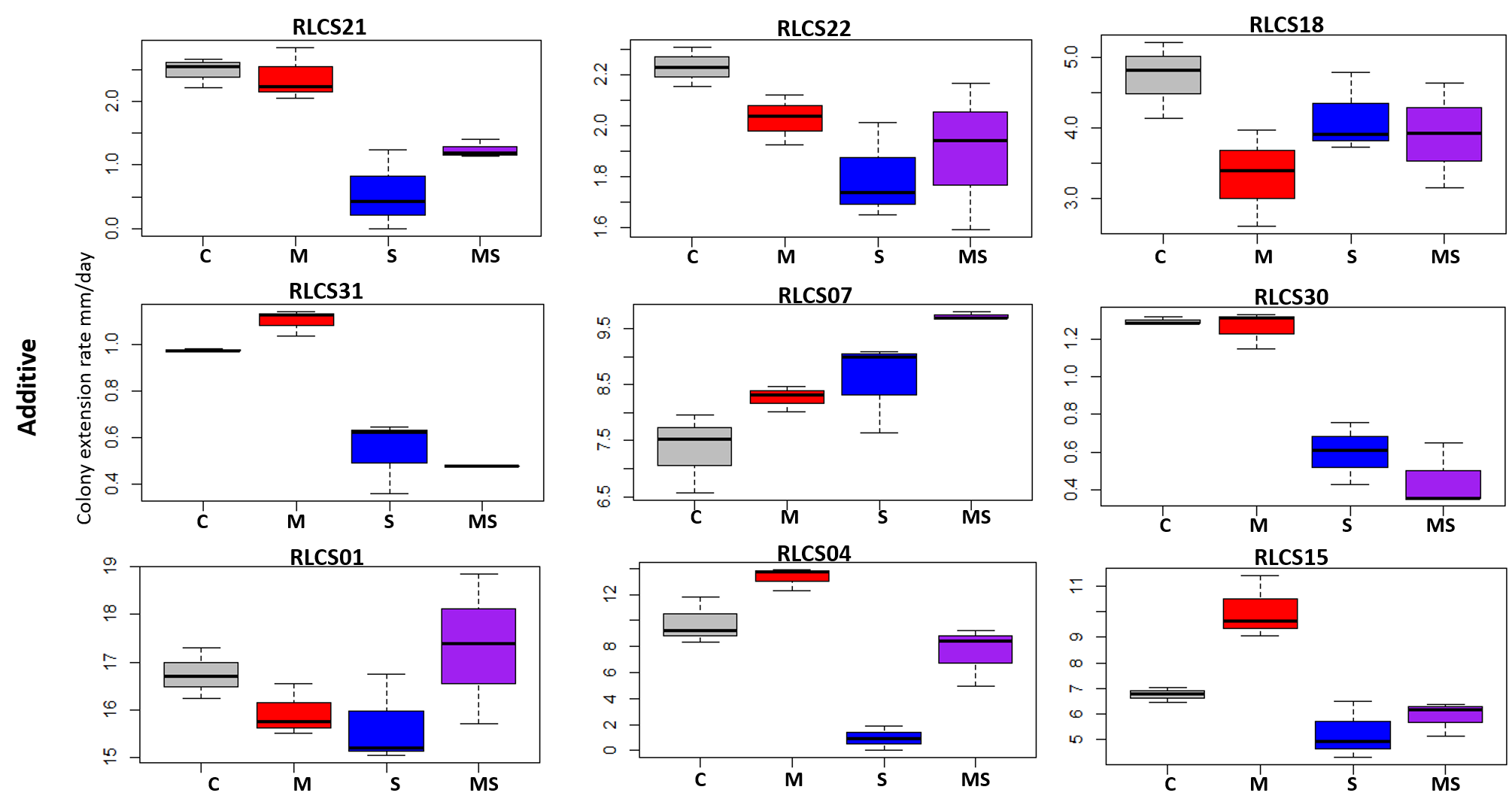

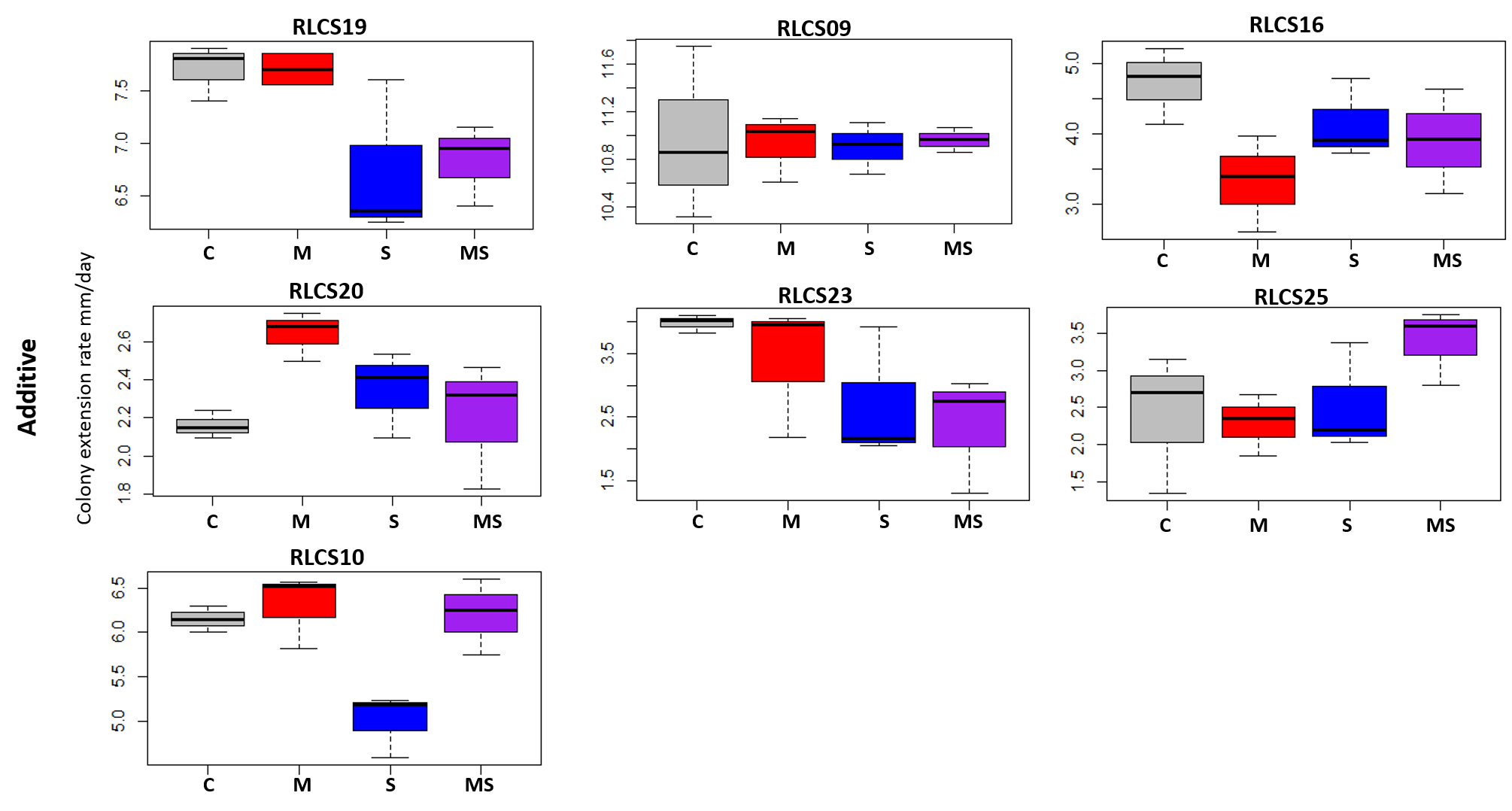
